# R2G2: A Python-R Framework for Seamless Integration of R/Bioconductor Tools into Galaxy

**DOI:** 10.64898/2025.12.22.695980

**Authors:** Jayadev Joshi, Fabio Cumbo, Daniel Blankenberg

**Affiliations:** Center for Computational Life Sciences, Cleveland Clinic Research, Cleveland Clinic, Cleveland, OH, USA; Department of Molecular Medicine, Cleveland Clinic Lerner College of Medicine, Case Western Reserve University, Cleveland, OH, USA

## Abstract

R is widely used in statistical computing, data analysis, and bioinformatics. A key contributor to its success in bioinformatics and computational biology is the open-source project Bioconductor. As of its latest release (3.20), the Bioconductor community offers 2,289 software packages for biomedical research, including genomic, transcriptomic, and proteomic analyses. Given R’s growing importance, integrating R and Bioconductor tools into platforms like Galaxy enhances accessibility, reproducibility, and scalability in bioinformatics workflows. The Galaxy Toolshed provides multiple tools leveraging R and Bioconductor packages. Additionally, various open-source public Galaxy servers, such as usegalaxy.org and usegalaxy.eu, already host several R and Bioconductor-based tools, highlighting the importance of past integration efforts. However, given the vast number of available packages, the full potential of R and Bioconductor within the Galaxy ecosystem remains underutilized. Galaxy’s web-based interface makes these powerful tools more accessible to researchers without programming expertise, fostering broader collaboration. Despite its advantages, integrating R packages into Galaxy can be complex. It requires XML wrappers to define inputs, outputs, and parameters, which can be time-consuming. Managing dependencies from CRAN and Bioconductor, resolving installation issues, and ensuring compatibility across different package versions further complicates the process. Many tools also require custom scripting, creating a steep learning curve for non-programmers. To address these challenges, we have developed a tool that automates the generation of Galaxy wrappers for R packages. This eliminates the need for manual XML writing, reduces complexity, and saves time. Our tool provides an intuitive interface for creating Galaxy-compatible tools without programming expertise and automates dependency management for seamless execution. Bioconductor has revolutionized bioinformatics, with thousands of researchers relying on its tools. Automating its integration into Galaxy removes technical barriers, democratizing access to advanced bioinformatics tools and workflows. Our solution bridges the gap between R-based analysis and user-friendly, scalable tools, ultimately advancing research accessibility and scientific discovery.

## 1. Introduction

In recent decades, the world has experienced unprecedented growth in data, particularly in the biomedical sciences, where high-throughput experiments and large-scale studies generate massive datasets. Community-driven initiatives, such as open-source software platforms and collaborative databases, have played a crucial role in enabling researchers to manage, share, and analyze these massive datasets effectively. (Silva et al. 2025; Mansueto et al. 2024). Programming languages, specifically R and Python, have played a crucial role in biomedical research by enabling rapid development of tools and algorithms to handle and analyze these massive and diverse datasets (Giorgi et al. 2022). In the recent past, we have observed an enormous growth in software packages related to these programming languages. For example, Bioconductor, the leading repository for R-based bioinformatics tools, has grown substantially and currently hosts 2,289 software packages as of 3.20 release, with approximately 75 new packages added annually through two releases each year. These include 928 annotation packages and 431 experiment data packages, supporting diverse genomic analyses. Similarly, Python’s bioinformatics ecosystem, exemplified by the Biopython project, benefits from the broader Python Package Index (PyPI), which contains over 614,000 packages as of March 2025. (Giorgi et al. 2022; Chan 2018; Staples 2023). Due to its math oriented community, R has become one of the most widely adopted programming environments when it comes to statistical computing and mathematics application in bioinformatics. Its active development community and rigorous standards for software interoperability and reproducibility have made R and Bioconductor an indispensable resource in the field (Siraji and Rahman 2023; Gentleman et al. 2004; Gentleman et al. 2005). Despite the availability of numerous powerful methods provided free of cost through these programming libraries, lack of advanced computational literacy among biologists remains the biggest limiting factor in the widespread adoption of these techniques. A similar trend is indicated by recent surveys that while bioinformatics tools are becoming essential in life science research, many researchers lack the confidence and training to use them effectively. In a Saudi Arabia-based survey of 309 scientists, 42.4% reported using bioinformatics tools in their research, but only 30.1% identified as working in bioinformatics-related fields. Among those using bioinformatics tools, more than half (51.9%) did so only occasionally. Furthermore, 56.4% of respondents acknowledged lacking sufficient bioinformatics knowledge (Alomair and Abolfotouh 2023). A global training survey by SEB/GOBLET reported that 57% of wet-lab scientists lacked confidence in using bioinformatics tools, 74% had no programming experience, and 58% felt uncomfortable with statistical methods (Attwood et al. 2019; Williams et al. 2019). These findings highlight a substantial gap between the growing demand for bioinformatics analysis and the practical ability of many researchers to perform such analyses independently (Balamurugan et al. 2021; Wilson Sayres et al. 2018; Verli and de Melo Minardi 2022). In recent years, open-source platforms such as Galaxy have played a significant role in bridging this gap by providing user-friendly, web-based graphical interfaces that help biologists adopt these advanced methods without computational expertise (Grüning et al. 2017; Joshi and Blankenberg 2022; Blankenberg et al. 2010). Integrating R and Bioconductor packages within Galaxy not only democratizes access to sophisticated statistical and bioinformatics methods but also facilitates collaboration across diverse research groups (Afgan et al. 2018; Blankenberg et al. 2010; Baichoo et al. 2018; Goecks et al. 2010; Langer et al. 2025). To date, 82 Bioconductor packages have been integrated with Galaxy; some of the most popular examples include DESeq2 (Love et al. 2014) and limma (Ritchie et al. 2015) for differential gene expression analysis. In a nutshell, developing Galaxy-compatible tools for R and Bioconductor packages typically requires writing XML-based tool wrappers to define inputs, outputs, and parameters, a process that is time-consuming, prone to error, and presents a steep learning curve for researchers without software development expertise (Cock et al. 2013; Joshi and Blankenberg 2022). Although the development of tools and workflows for Galaxy is largely community-driven, it still requires a substantial amount of human labor and time to develop, test, and maintain high-quality tools for the research community. In addition to this, managing dependencies from CRAN and Bioconductor, resolving package installation issues, and ensuring compatibility across different software versions introduce additional technical challenges.

To address these barriers, save time and labor, and reduce technical complexity, we have developed R2G2, a Python- and R-based package that streamlines the integration of R packages into the Galaxy platform. This solution eliminates the need for manual XML wrapper creation, simplifies dependency management, and provides an intuitive interface for generating Galaxy-compatible tools. By automating these tasks, R2G2 lowers technical barriers, accelerates tool development, and broadens access to R and Bioconductor’s extensive resources as thousands of investigators worldwide relying on their tools for advanced biological data analysis (Ruprecht et al. 2024; Giorgi et al. 2022). By automating the integration of R and Bioconductor packages into Galaxy, R2G2 democratizes access to these resources, helps bridge the computational skills gap identified by recent surveys, and fosters broader collaboration between computational and experimental scientists. This work contributes to ongoing community-driven efforts to make state-of-the-art bioinformatics tools more accessible, scalable, and reproducible for the wider life science research community.

## 2. Material and Methods

### 2.1 The main motivation of the work

The primary objective of this project was to develop a comprehensive framework to facilitate the integration of valuable R and Bioconductor functionalities into Galaxy. While the R and Bioconductor ecosystem offers an extensive collection of R-based packages for statistical analysis and visualization of various omics datasets, its adoption within integrative workflow environments such as Galaxy remains limited due to the previously mentioned challenges. To achieve this, we designed a Python and R-based library that automates the generation of wrapper scripts for Bioconductor packages, thereby enabling their seamless deployment as Galaxy tools. Python was selected as the implementation language because of its interoperability, wide adoption in the bioinformatics community, and natural compatibility with the Galaxy ecosystem.

The library systematically manages package dependency resolution, argument parsing, and input/output standardization, thereby reducing the burden on developers who would otherwise need to manually configure wrappers. Beyond building wrappers directly from CLI (command-line interface) architecture based R scripts, we also implemented a dedicated module capable of generating Galaxy wrappers for complete R packages. In both approaches, the workflow begins with either an R script or a Bioconductor package. The subsequent step involves identifying and mapping the arguments into Galaxy input and output parameters. Finally, the system generates the command-line interface along with the input and output sections required for a fully functional Galaxy wrapper. The full implementation details are as follows. A complete framework has been shown in Figure 1:

**Figure 1.**
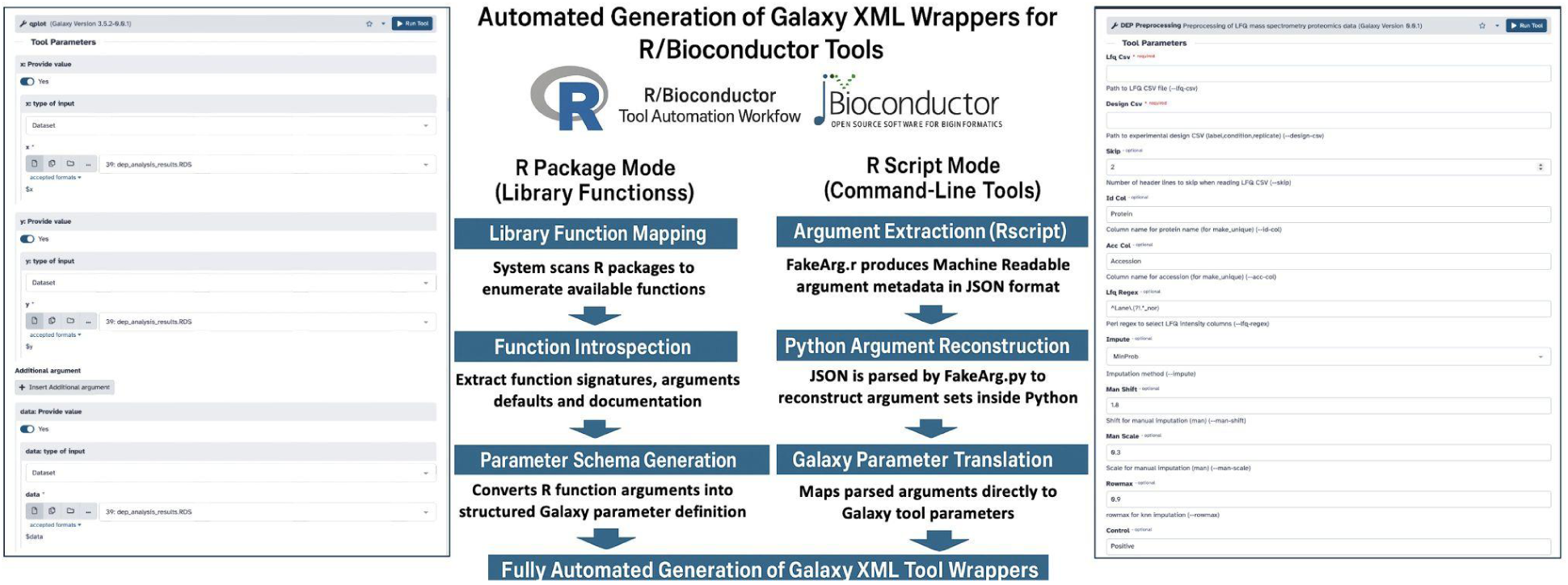
Automatic Generation of R -based tools to integrate into the Galaxy platform. This workflow illustrates how R library functions are mapped and converted into Galaxy tools. On the left, R library packages are parsed to extract function objects, arguments, and definitions, which are then translated into Galaxy parameter sets. On the right, from a R command line script, FakeArg.r script exported arguments in the python format as JSON objects, which are converted back into Python code via a custom argument parsing class “CustomFakeArg”. These arguments are subsequently mapped to Galaxy parameters. Together, both processes enable the automatic generation of XML wrappers to integrate R-based bioinformatics tools within Galaxy’s graphical user interface (GUI). The GUI on the right side was created using the *ggplot2* library, while the GUI on the left side was generated using an Rscript based on the *DEP* package.

### 2.2 System Architecture and Environment

A detailed overview of the system architecture is as follows

#### 2.2.1 Galaxy Platform Setup

The R2G2 library can generate XML wrappers that are compatible with the Galaxy software and function across multiple Galaxy releases. Additionally, the provided tool template allows flexibility to specify a particular tool profile when needed. Additionally, the provided tool template allows flexibility to specify a particular tool profile when needed.

#### 2.2.2 R Version and Bioconductor Release

By default, Galaxy tools rely on Conda environments for dependency resolution and reproducibility. Therefore, when generating a Galaxy tool wrapper from an R package using the R2G2 library, it is essential that a corresponding Conda dependency is available. This ensures that the required R version, Bioconductor release, and associated packages can be correctly installed and executed within Galaxy. Similarly, when generating wrappers from R scripts, the underlying R package should also be available as a R or Bioconductor Conda package. This integration allows R2G2 to seamlessly reference the correct dependencies and manage them through Conda (or alternatives such as mamba, Docker, or Singularity when needed), ensuring consistent runtime environments across Galaxy instances.

#### 2.2.3 Generating Galaxy tools from R packages

Developing software packages for various methods and algorithms is one of the most common and effective approaches for disseminating code within the scientific community. This practice enhances reproducibility and facilitates the broader adoption of new methodologies. By providing ready-to-use functions and classes, packages enable researchers to utilize existing algorithms without the need to reinvent the wheel, while also supporting the development of novel algorithms that can build upon established functionalities. Given the broad availability of R and Bioconductor packages, we implemented functionality to automatically generate Galaxy wrappers directly from R packages. The implementation details are as follows:

##### 2.2.3.1 Wrapper Generation Workflow

The Python script “r2g2_on_package.py” automatically converts R library functions into Galaxy-compatible tool wrappers. The script leverages the rpy2 interface to interact with R packages, extract function signatures, and dynamically generate the corresponding Galaxy tool XML definitions. This process ensures that R-based functionality can be seamlessly integrated into Galaxy workflows without requiring extensive manual tool wrapper development.

##### 2.2.3.2 Input Parameters

The script accepts several command-line arguments to generate Galaxy tool wrappers. The “--name” parameter specifies the R package to be wrapped and is required. The “--package_name” argument defines the corresponding Conda package name, which is optional and defaults to the R package name if not provided. Similarly, the ‘--package_version” argument allows the user to specify the Conda package version, with the default set to the detected version. The “--out” parameter designates the output directory for the generated Galaxy wrapper files, which defaults to out/. Finally, the “--galaxy_tool_version” argument assigns a version string to the Galaxy tools, with a default value of 0.0.1. Full parameter details are integrated with the R2G2’s “--help” command.

##### 2.2.3.3 R Package Import and Metadata Extraction

We utilized the rpy2 package to enable seamless integration of R and Bioconductor based tool Generation. The rpy2.robjects.packages module acts as Python’s gateway to the R package ecosystem, with its core function importr() calling R’s library() and wrapping the resulting R namespace into a Package object. This makes R functions accessible as callable Python objects through SignatureTranslatedFunction. R2G2 leverages these capabilities to extract metadata and other R objects, dynamically importing the specified R package. The package version is automatically detected unless explicitly provided. Based on this information, a macro XML file is generated, containing reusable tool components such as requirements, macros, and versioning details.

##### 2.2.3.4 Function Iteration and Documentation

The next step is to iterate over each function in the imported R package. For each function:

a. **Function Discovery:** Functions are identified via dir() on the package object.
b. **Documentation Retrieval:** Help pages are accessed through rpy2.robjects.help.pages, which extracts the underlying R help files (.Rd documentation associated with the R object). When available, these .Rd entries are converted to reStructuredText (RST) and embedded into the tool’s help section. If multiple help pages exist, all are concatenated. When no R documentation is found, the Python docstring is used as a fallback.
c. **Tool Metadata Initialization:** A metadata dictionary is constructed, containing the Galaxy tool ID, version, function name, description, and other required XML fields.

##### 2.2.3.5 Parameter Processing

Function parameters are analyzed using the package_obj.formals() method, which retrieves the formal arguments defined for each R function. For every parameter, the script identifies its default value, data type (e.g., integer, floating point, string, or logical/boolean), and whether it represents a single or multiple input. Each parameter is then mapped to the corresponding Galaxy XML input template (Table 1), enabling seamless integration of function arguments into the Galaxy tool interface. When the parameter type cannot be reliably inferred, a generic not_determined template is applied to ensure flexibility in handling diverse input types. Furthermore, the script incorporates specialized handling for the ellipsis (…) parameter, which is converted into repeatable and conditional Galaxy inputs, thereby supporting functions that accept variable-length argument lists.

**Table 1.** Mapping of R argument parsing parameters to corresponding Galaxy tool input and output parameters. The table illustrates how commonly used R command-line arguments (defined via argument parsers) can be systematically translated into Galaxy XML tool definitions, enabling seamless integration of R-based tools within the Galaxy platform.

| **R Argument Parsing (argparse)** | **Meaning in CLI** | **Galaxy Tool Parameter** | **Notes / Translation Rule** |
| --- | --- | --- | --- |
| type = "character" | String input | <param type="text"> | Default text input in Galaxy. |
| type = "integer" | Integer input | <param type="integer"> | Restrict input to integers. |
| type = "double" / "numeric" | Floating-point input | <param type="float"> | Maps to numeric entry. |
| action = "store_true" | Boolean flag (set True if present) | <param type="boolean" truevalue="--flag" falsevalue="" /> | Common for switches like --verbose. |
| action = "store_false" | Boolean flag (set False if present) | <param type="boolean" truevalue="" falsevalue="--flag" /> | Inverted flag logic. |
| choices = c("A","B","C") | Restrict to set of values | <param type="select"> with <options> | Drop-down menus in Galaxy. |
| nargs = "+" or "*" | Multiple values allowed | <param type="data" multiple="true" /> | <repeat> |
| metavar = "FILE" | Expected input file | <param type="data"> | Galaxy handles file input. |
| Subparsers/Mutually exclusive groups | Argument parsing groups | <conditional> | Galaxy conditional parameter block |

##### 2.2.3.6 R Script Generation

For each wrapped function, an R script block is programmatically generated. This script begins by loading the required R package [library(package_name)], then maps Galaxy inputs to the corresponding R function arguments with appropriate type conversions, such as using readRDS for datasets or applying string quoting for text values. Once the inputs are processed, the script executes the R function with the mapped arguments and saves the resulting object(s) in RDS format, a native binary file format in R (R Data Serialization) used to store single R objects and later read them back into an R session, ensuring compatibility with downstream Galaxy tools. However, RDS format relies on R’s internal serialization mechanism, therefore it shares the same security drawbacks as Python’s pickle format. The bigger security drawback is that the deserialization process can execute embedded code. In practice, this risk can be avoided by restricting file sources to controlled environments and using alternative portable formats for data sharing.

##### 2.2.3.7 XML Tool Wrapper Creation

The processed metadata, function documentation, parameter specifications, and generated R script are inserted into an XML tool template (tool_xml). Each function results in a corresponding Galaxy tool XML file saved in the output directory. The tool IDs are automatically sanitized (removing invalid characters) to meet Galaxy ToolShed requirements. Optionally, a specialized r_load_matrix.xml wrapper is generated to facilitate matrix-to-RDS conversion. This wrapper serves as a useful helper tool that allows Galaxy users to seamlessly transform tabular data into a format that R functions, and the automatically generated wrappers, can readily interpret and process. By bridging the format gap between Galaxy’s standard tabular inputs and R’s native RDS format, it ensures compatibility, since most R functions require structured R objects rather than raw text. Moreover, it standardizes inputs by providing a consistent mechanism for reading data once converted into RDS, thereby simplifying integration and enhancing interoperability across all wrapped R functions.

##### 2.2.3.8 Error Handling

The script includes robust error handling. If a function cannot be processed (e.g., due to undocumented parameters or unsupported argument types), the function is skipped with a logged warning.

#### 2.2.4 Generating Galaxy tools from R scripts

Another common and user-friendly way of distributing algorithms is through command-line tools implemented in programming languages. Libraries such as argparse in Python and R make command-line argument handling more intuitive and powerful. These argument parsing based scripts enable the creation of more comprehensive and complete tools which require almost no adjustment and can be used directly under a Galaxy tool. A detailed description is provided below, outlining the in-depth workflow of this implementation

##### 2.2.4.1 R Script Wrapper Generation

To enable Galaxy tool integration for R scripts with command-line arguments, we implemented a Python-based wrapper generation workflow. This functionality enables users to generate Galaxy tool wrapper for an Rscript that can run as a command line tool implemented based on the r-argparse library.

##### 2.2.4.2 Converting R based arguments to Python based arguments

The first step in this process is to extract information about the command-line arguments that an R script accepts. To achieve this, we implemented the FakeArg.r script based on the r-argument package. The FakeArgs class is implemented using the R6 package in R, which provides an object-oriented programming (OOP) system. This OOP system offers a reference-based approach and supports a traditional object-oriented programming style similar to that of Python, Java, or C++, including features such as methods, fields, inheritance, and encapsulation. The FakeArgs class leverages the native functionality of the r-argparse library to extract arguments from the R script and convert them into Python-compatible strings. These Python-compatible strings, which contain the argument definitions, are then saved into a JSON file for downstream processing.

##### 2.2.4.3 Convert arguments strings into Python code

In the next step, we utilized json_to_python, json_to_python_for_param_info, and extract_simple_parser_info functions to assists in categorizing whether an argument belongs to a conditional block. This is determined by iterating over the arguments and inspecting their grouping, such as subparsers or mutually exclusive groups.

##### 2.2.4.4 Dynamically extracting the parameters from the converted Python arguments

In this step, we leverage the anvi’o (Eren et al. 2020) tool-wrapper-generation package. Specifically, our CustomFakeArg class extends the functionality of the FakeArg class provided by the anvio package (from anvio import FakeArg), which itself inherits from Python’s standard argparse library, thereby enabling structured handling of command-line arguments within the wrapper generation workflow. Parameter metadata, including names, types, and categories, is extracted using CustomFakeArg. This information is then encapsulated within a blankenberg_parameters object, which provides methods to generate conditional input blocks, mutually exclusive groups, miscellaneous parameters, and corresponding command-line representations. These components are subsequently combined to construct the full tool command section, ensuring that the original R script can be executed via Galaxy with correctly mapped arguments

##### 2.2.4.5 Mapping Extracted parameters to Galaxy tool wrapper

Once all parameter details, including names, types, metadata, and information about various groups such as mutually exclusive groups or subparsers, are obtained, the next step is to convert them into the corresponding components of a Galaxy wrapper. Various methods of the CustomFakeArg class, such as generate_conditional_block, generate_mutual_group_conditionals, generate_misc_params, and generate_command_section_subpro, are used to construct individual sections of the tool.

##### 2.2.4.6 Dependencies management

Dependencies required by the R script are identified using return_dependencies function and formatted into Galaxy-compatible <REQUIREMENTS= tags via return_galax_tag. This ensures that all necessary R packages are available in the Conda environment at runtime.

##### 2.2.4.7 XML Wrapper Generation

All components generated in the previous steps, including input blocks, output blocks, dependency blocks, and command-line sections, are fed into a Jinja2-based XML template (from jinja2 import Template) to generate the complete Galaxy XML wrapper. The tool metadata, such as ID, name, version, description, inputs, outputs, commands, help text, and dependencies is incorporated into the template to produce a robust and fully defined Galaxy tool. The resulting XML file is written to the specified output directory, and any temporary working directories and intermediate files are removed to maintain a clean workspace. This workflow enables any R script with argument parsing to be automatically converted into a fully functional Galaxy tool wrapper, bridging the gap between standalone R scripts and reproducible Galaxy workflows while ensuring consistent parameter handling, dependency management, and standardized input/output formats.

#### 2.2.5 Integration with Galaxy Toolshed

The Galaxy ToolShed is a central repository for sharing Galaxy tool wrappers, enabling researchers to easily access, install, and use a wide range of tools within the Galaxy platform. It serves as both a distribution hub and a version-controlled archive, helping maintain consistent tool functionality across different Galaxy instances. To facilitate the development and deployment of high-quality tools, the Planemo toolkit can be used. Planemo provides commands to lint, test, and validate Galaxy tool wrappers locally before submission, ensuring adherence to Galaxy’s standards and best practices. Once a wrapper passes validation, Planemo can be used to push it directly to a Galaxy ToolShed repository, streamlining the process of distribution and making the tool readily available to the broader Galaxy community. By leveraging ToolShed and Planemo, developers can maintain reliable, reproducible tools while simplifying installation and sharing across the broader Galaxy and research community.

#### 2.2.6 Reproducibility

For reproducibility, Galaxy supports containerization via Docker and Singularity, allowing users to deploy pre-installed Galaxy instances with the generated R-based tools, along with supporting data libraries and workflow suites. This approach enables researchers to provide a fully configured, containerized environment that ensures a robust and reproducible data analysis experience with the developed tools. These workflows not only provide access to the tools but also guarantee reproducibility of analyses, making them available to a global community of over 600,000 Galaxy users. By combining containerized Galaxy instances, ToolShed distribution, and workflow sharing, developers can deliver reliable, reproducible, and easily accessible computational tools and workflows at scale.

## 3. Results and Discussion

To evaluate the effectiveness of our automated Galaxy wrapper generation tool, we focused on integrating R and Bioconductor packages into the Galaxy platform. Given the extensive repertoire of over 2,200 Bioconductor packages, manual wrapper creation is time-consuming and prone to errors, especially for complex tools with multiple inputs, outputs, and interdependent parameters. Using our approach, wrappers can be generated automatically from R scripts, significantly reducing development time while ensuring proper handling of dependencies and compatibility across versions. We applied the tool to both in-house and open-source R scripts, including widely used packages such as ggplot2, demonstrating its capability to produce fully functional Galaxy tools. This automation not only streamlines the integration process but also enhances reproducibility, accessibility, and scalability, enabling researchers without programming expertise to leverage advanced R-based analyses within Galaxy workflows. By bridging the gap between Bioconductor and Galaxy, our approach facilitates the broader adoption of R-based bioinformatics tools, supporting more reproducible and collaborative research practices.

We present several use cases leveraging publicly available R scripts alongside in-house developed R scripts for various biological analyses

### 3.3.1 Generating wrappers from R scripts that support command-line argument parsing

#### 3.3.1.1 Based on in-house generated R scripts

Integrated analysis toolset for robust and reproducible analysis of mass spectrometry proteomics data.

a. **Dataset** In this use case, we utilized the processed example dataset provided by the DEP package (Zhang et al. 2018). The original data belong to the authors of the study in which ubiquitin-protein interactors were characterized (Zhang et al. 2017). Before supplying the data with the DEP package, the raw mass spectrometry data was processed using MaxQuant (Zhang et al. 2017; Cox and Mann 2008), and we simply utilized the resulting dataset provided through the DEP package. We used this publicly available processed file exclusively to demonstrate the usability of R2G2 on our in-house generated R script for automated Galaxy wrapper generation.
b. **DEP_preprocessing.r CLI tool** enables automated preprocessing and quality control of label-free quantitative proteomics data within Galaxy. Implemented command-line R module wraps the core DEP preprocessing functions behind a structured argument interface. This script is providing reproducible and parameterized execution of all major preprocessing steps, including filtering, normalization, imputation, and QC visualization, Figure S1. The tool takes two primary inputs: the unique proteins table with LFQ (Label Free Quantification) intensity values and the experimental design file, both supplied in CSV format. Users may also specify the prefix used for LFQ intensity columns, allowing compatibility with datasets from different quantification pipelines (e.g., MaxQuant, FragPipe). Using these inputs, the script constructs the SummarizedExperiment object required for DEP-based processing. Following normalization, protein intensities are imputed using the user-selected method (e.g., MinProb, MinDet, kNN, QRILC, MLE, bpca). The resulting imputed dataset is saved in both RDS and CSV formats for downstream analysis. Additional diagnostic plots, including imputation density plots, are generated to help users evaluate the effect of the imputation strategy. For convenience, the tool produces a combined PDF containing all generated plots, facilitating rapid review of preprocessing quality in a single document. Overall, the generated Galaxy wrapper based on this R script, with the help of R2G2 package, provides a comprehensive and flexible preprocessing pipeline for LFQ proteomics data within Galaxy, enabling users to perform standardized QC, normalization, and imputation without requiring direct interaction with R.
c. **DEP_DE_analysis.r CLI tool** exposes the core DEP differential expression and visualization functions through a standardized argument interface. The tool accepts an imputed DEP SummarizedExperiment object (RDS format) and performs differential expression testing using test_diff() based on a user-selected control condition, Figure S2. Significance thresholds for P-value (α) and log₂ fold change can be set through Galaxy parameters, and the tool automatically annotates significant proteins using add_rejections(). All differential expression results are exported as a CSV file for downstream use within Galaxy workflows. In addition, the script provides two plot-generation modes, PCA and volcano, implemented based on the subparsers. Users can configure principal components, the number of variable proteins, point size, contrast definitions, label size, and whether to display protein names. The script generates publication-quality plots using the corresponding DEP functions (plot_pca() or plot_volcano()) and saves them to the Galaxy-designated output directory.

Overall, the generated Galaxy wrapper based on this R script with the help of R2G2 package serves as the computational backend for the Galaxy tool, enabling flexible, reproducible execution of DEP differential expression analysis and visualization directly within Galaxy’s workflow system. It provides parameterized control, standardized outputs, and seamless integration with upstream preprocessing and downstream interpretation tools.

#### 3.3.1.2 Based on open-source R scripts

To demonstrate the usability and versatility of the R2G2 package, we downloaded publicly available R script-based tools implemented in a command-line style using the r-argparse library. In total, we collected around 41 different tools, some of which are highlighted below, with details presented in Table 2.

**Table 2.** Summary of publicly available R scripts for computational biology analysis, used in this study to demonstrate the usability of R2G2 in automatic Galaxy tool generation.

| S. No. | Script name | Category |
| --- | --- | --- |
| 1 | indelfindr.R | Genetic variants detection |
| 2 | CpG_island_identificator.R | CpG Island research |
| 3 | DESeq2.R | DE analysis |
| 4 | edgeR.R |  |
| 5 | extract_or_remove_and_split_from_multifasta.R | Fasta file manipulation for sequence analysis |
| 6 | extract_or_remove_seqs_from_fasta.R |  |
| 7 | fasta_select_or_remove_by_header_pattern.R |  |
| 8 | fasta_split_to_multifasta_by_win_step.R |  |
| 9 | Update fasta_split_to_multifasta_by_win_step.R |  |
| 10 | fasta_split_to_singlefastas_by_win_step.R |  |
| 11 | Update fasta_split_to_singlefastas_by_win_step.R |  |
| 12 | merge_multifasta_to_singlefasta.R |  |
| 13 | merge_seqs_from_singlefasta.R |  |
| 14 | parse_headers_from_fasta.R |  |
| 15 | select_fasta_or_header_by_length.R |  |
| 16 | singlefasta_to_multifasta.R |  |
| 17 | split_multifasta_to_multifastas.R |  |
| 18 | split_multifasta_to_singlefasta.R |  |
| 19 | trim_multifasta.R |  |
| 20 | combine_dssp_statistics_to_table.R | PDB data analysis |
| 21 | pdb_to_fasta.R |  |
| 22 | split_pdb_to_chains.R |  |
| 23 | split_pdb_to_fasta.R |  |
| 24 | trim_pdb.R |  |
| 25 | trim_pdb_by_chain.R |  |
| 26 | trim_pdb_to_fasta.R |  |
| 27 | Update trim_pdb_to_fasta.R |  |
| 28 | trim_pdb_to_fasta_by_chain.R |  |
| 29 | Update trim_pdb_to_fasta_by_chain.R |  |
| 30 | all_aas_content_multifasta.R | Sequence analysis and statics |
| 31 | calculate_and_select_aa_content.R |  |
| 32 | calculate_and_select_at_content.R |  |
| 33 | calculate_and_select_by_length.R |  |
| 34 | calculate_and_select_gc_content.R |  |
| 35 | calculate_biophysical_properties_multifasta.R |  |
| 36 | get_length_from_multifasta.R |  |
| 37 | multifastas_gc_at_stats.R |  |
| 38 | multifastas_length_stats.R |  |
| 39 | select_fasta_or_header_by_aa.R |  |
| 40 | select_fasta_or_header_by_at.R |  |
| 41 | select_fasta_or_header_by_gc.R |  |

a. **INDELfindR** is an open-source command-line tool developed in R for the detection of both simple and complex insertion-deletion (INDEL) variants, a common form of genetic variation where nucleotides are either inserted into or deleted from the genome. INDELs can have profound biological consequences, often altering coding sequences, disrupting protein function, or affecting gene regulation. Detecting these variants is therefore essential for understanding genetic diversity, disease mechanisms, and potential biomarkers. The tool processes sequencing data and outputs INDEL calls in VCF v4.3 format, making the results directly compatible with widely used downstream analysis and annotation pipelines. This ensures smooth integration with existing workflows for variant annotation, population genetics studies, and clinical interpretation. With INDELfindR, researchers can analyze data from whole-genome or exome sequencing experiments to identify INDELs linked to disease-associated genes, functional variants that impact protein structure, or candidate markers for diagnostics and therapeutic targeting. By providing a streamlined, reproducible, and standards-compliant workflow, INDELfindR supports both fundamental research and translational applications in genomics and precision medicine (https://github.com/TranslationalBioinformaticsLab/INDELfindR).
b. **biomisc_R** biomisc_R is a collection of command-line bioinformatics scripts written in R. This toolkit provides modular utilities for common tasks such as handling FASTA files, analyzing sequence statistics, identifying CpG islands, performing differential-expression analysis, manipulating PDB structures, and processing synthetic biology constructs. The repository leverages essential R packages including r-argparse, DESeq2, edgeR, ape, phylotools, stringr, bio3d, Biostrings, GeneGA, and Peptides to achieve functionality across domains like structural bioinformatics, genomics, and statistics (https://github.com/olgatsiouri1996/biomisc_R).

#### 3.3.1.3 Rscripts from Bioconductor packages

In this study, we systematically collected and analyzed 2,289 Bioconductor packages to extract Rscripts that utilize the argparse library for command-line argument parsing. From these packages, we identified 51 Rscripts, reported in the Table 3, belonging to key tools, including *CircSeqAlignTk*, *MAGAR*, *RnBeads*, *infercnv*, and *openCyto*. Using R2G2, Galaxy wrappers for these Rscripts were generated and are provided alongside this manuscript. This approach demonstrates that automated wrapper generation can standardize script execution and simplify integration into reproducible pipelines. The wrappers enable streamlined access to diverse analytical functions, reduce manual intervention, and enhance scalability across multiple datasets. Overall, this work illustrates the feasibility of creating a comprehensive, command-line-based interface for a wide array of Bioconductor tools, supporting more efficient and reproducible computational analyses in genomics and proteomics research.

**Table 3.**
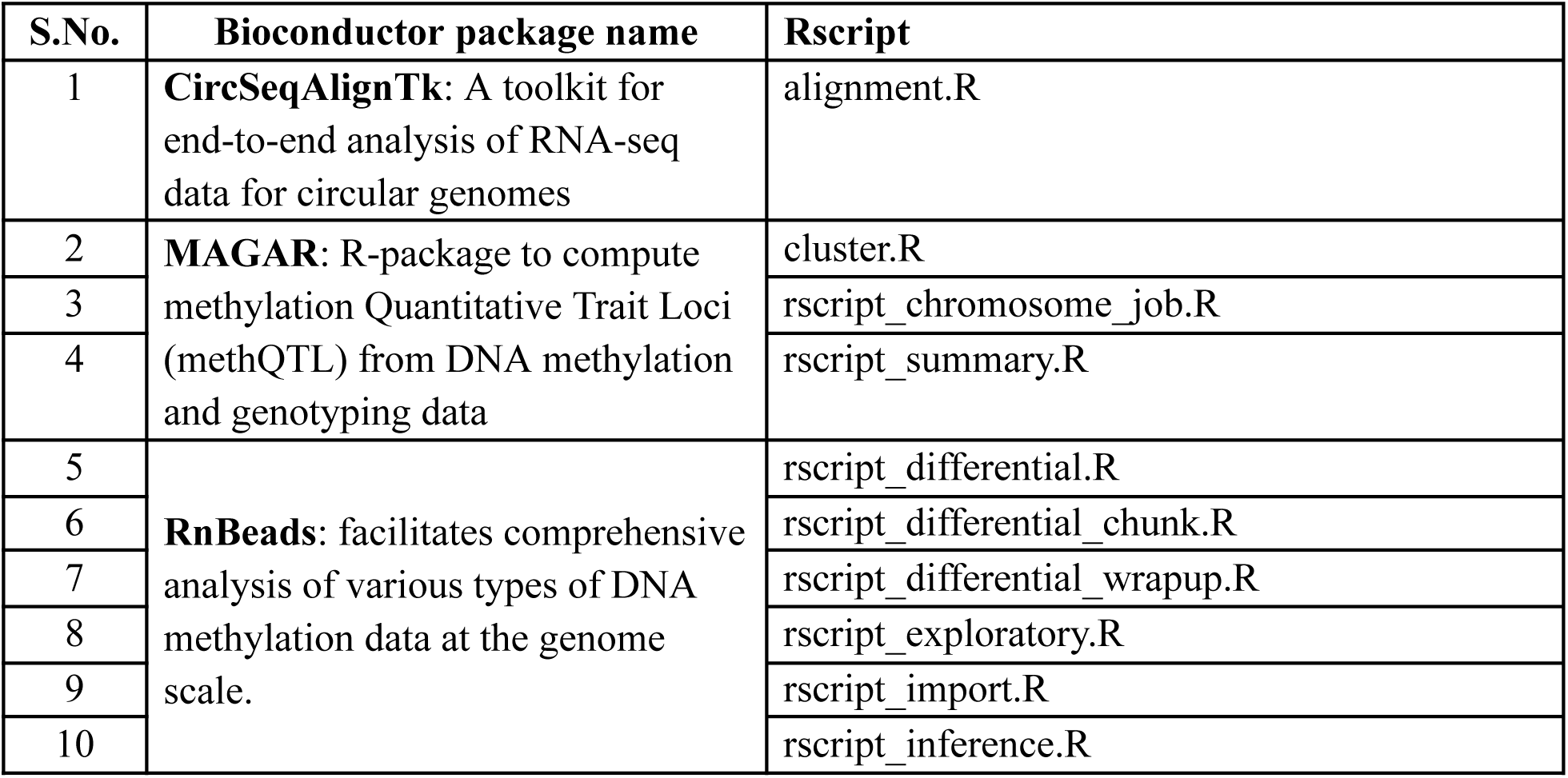
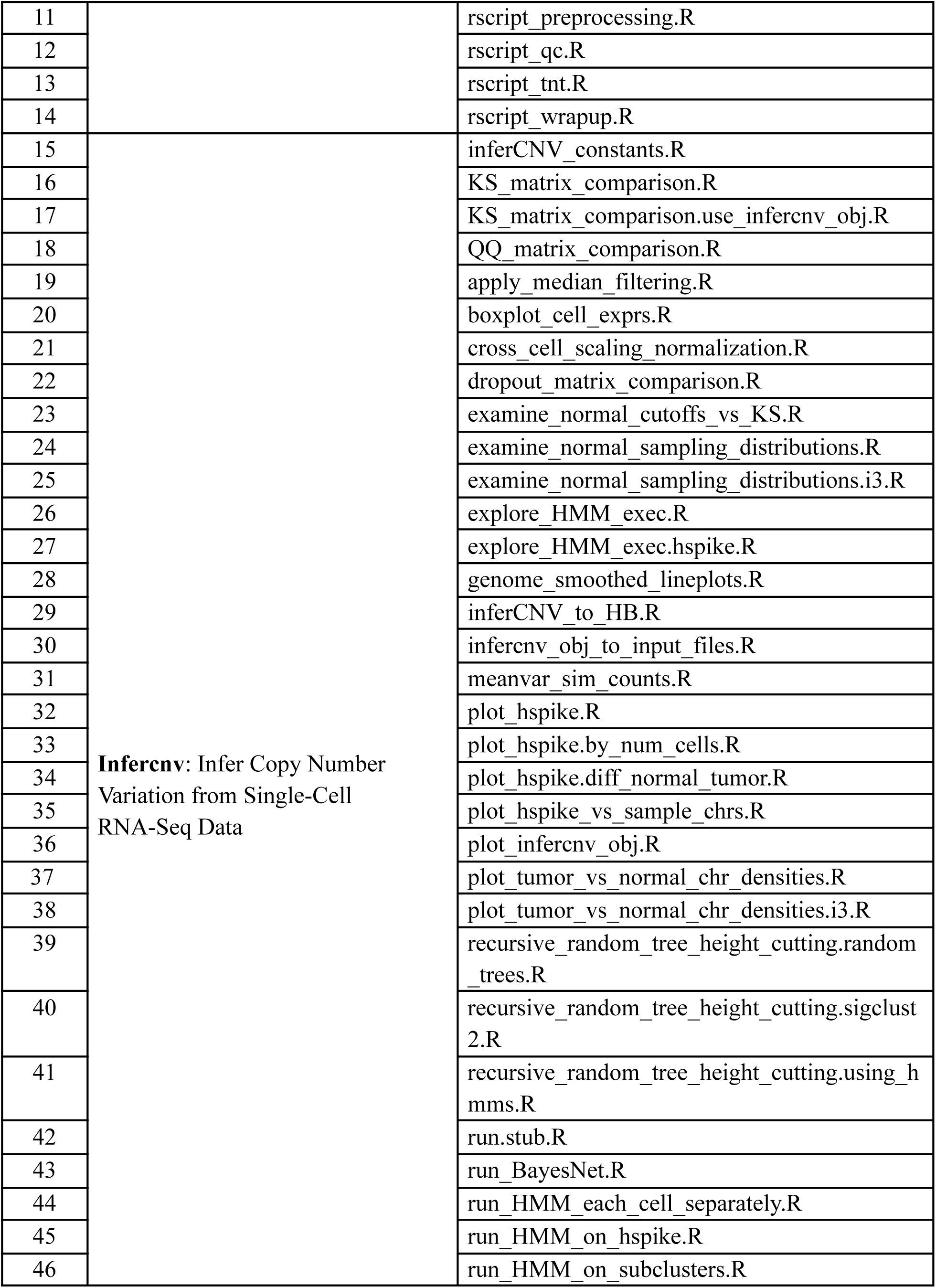
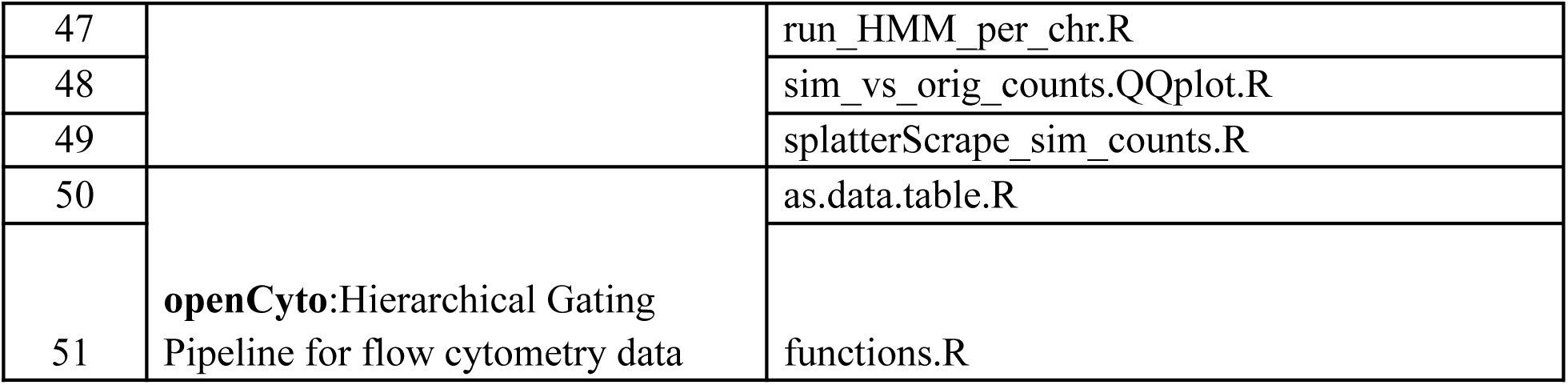
Summary of the Bioconductor packages and Rscripts extracted to demonstrate the usability of R2G2 in automatic Galaxy tool wrapper generation.

### 3.3.2 Generating tool wrappers from R package

The ggplot2 package is one of the most widely used libraries in R for creating high-quality, publication-ready visualizations. By generating Galaxy tool wrappers for ggplot2, we demonstrate how standard visualization tasks can be seamlessly integrated into the Galaxy platform. This use case highlights the automation of wrapper creation for the core functions of the ggplot2 library, enabling users to access advanced plotting capabilities and leverage these building blocks for developing more sophisticated Galaxy tools. R2G2 iterates over all available objects in the ggplot2 library, identifies functions based on their signatures, and determines their arguments to generate corresponding Galaxy tool parameters. Through this process, more than 450 Galaxy tool wrappers are created. While the usability of these wrappers as standalone tools is limited for comprehensive analyses, the primary motivation behind this implementation is to dynamically extract function-level information and provide a complete set of building blocks for robust Galaxy tool development. These extracted wrappers can then be combined to assemble more complex and practical tools, particularly in cases where argument-parsing scripts are not readily available.

Despite this, the extensive array of available R and Bioconductor packages indicates that their full potential within the Galaxy ecosystem has yet to be realized. The results demonstrate that the R2G2 package provides comprehensive and detailed support in streamlining the development of Galaxy tools for R-based packages. Efforts such as R2G2 are essential for harnessing the true power of these packages in bioinformatics analyses. By leveraging Galaxy’s web-based interface, these tools become more accessible to researchers without programming expertise, promoting broader collaboration and facilitating the adoption of advanced computational analyses across diverse research communities. R2G2 not only supports wrapper generation for R-based scripts implemented using argument parsing but also enables package-based wrapper generation, thereby enhancing usability and providing broader coverage in R-based tool development. Despite the powerful functionality of the R2G2 package, it is reasonable to highlight some limitations we observed during testing, which could inform future development and enhancement of the package. Currently, R2G2 attempts to infer output parameters based on keywords, specifically by checking if an argument name begins with certain predefined terms such as “output.” While this heuristic can often identify output parameters, it has limitations and may fail to correctly distinguish true outputs from inputs.As a result, in some cases certain input and output parameters may be misassigned if the user relies entirely on the automatic input-output detection mechanism. This necessitates a brief manual review and adjustment after wrapper generation. To make output parameter definition more robust and straightforward, we have introduced an argument-based output parameter specification, which effectively addresses these scenarios. This highlighted the need to implement a comprehensive and robust automated testing framework for the tools generated by R2G2, allowing users to perform broad-coverage tests quickly and thereby improving tool reliability. Currently, we rely on the default testing approach, which limits the automated testing capabilities of R2G2. We aim to enhance and expand this functionality in future versions. Generating tool wrappers directly from packages can sometimes produce an enormous number of tools, not all of which are immediately useful. Users must identify the desired function-based tools and merge them into more practical and efficient workflows. While R2G2 simplifies this process, it still requires user intuition and creativity to make these individual wrappers truly useful, which can sometimes limit the overall usability of this approach. Nevertheless, this represents the first approach of its kind, and in future versions, we aim to make this functionality more robust, enabling the generation of more practical tools directly from packages, even when a command-line R script is not available. Overall, R2G2 not only helps new users learn and implement useful tools in Galaxy but also saves considerable effort in creating complex Galaxy tool wrappers. R2G2 can generate these complex wrappers in seconds with a single command, reducing substantial amounts of the human effort typically needed for developing such tools.

## Conclusion

In this study, we present a comprehensive framework for automatically generating Galaxy tool wrappers from R packages and scripts, with a focus on Bioconductor and ggplot2 workflows. By leveraging R2G2, we demonstrated how individual functions and argument parsing information can be dynamically extracted to create robust, reusable building blocks for Galaxy tools. This approach significantly reduces the time and technical expertise required to integrate R-based analyses into the Galaxy platform, while maintaining reproducibility, standardized input/output formats, and dependency management. Overall, the framework provides a scalable and flexible solution for democratizing access to R-based bioinformatics tools. Ultimately, this work bridges the gap between R’s extensive computational capabilities and Galaxy’s user-friendly interface, facilitating reproducible, accessible, and scalable data analysis for the broader research community.

## Availability

The R2G2 package can be installed either directly from PyPI or from its GitHub repository. For a simplified installation, R2G2 is available as a PyPI package (https://pypi.org/project/r2g2/0.1.1/) and can be installed using standard Python package managers. Alternatively, the source code can be downloaded directly from GitHub at https://github.com/BlankenbergLab/r2g2, where detailed installation and usage instructions are provided. All open-source R scripts used to demonstrate the usability of R2G2 were obtained from publicly available GitHub repositories, including https://github.com/TranslationalBioinformaticsLab/INDELfindR, https://github.com/olgatsiouri1996/biomisc_R, and the respective GitHub repositories of the referenced Bioconductor packages. All Galaxy-compatible R script wrappers generated in this work to demonstrate the capabilities of R2G2 are available at https://github.com/jaidevjoshi83/galaxy_tool_wrappers.

## Author Contributions

DB conceived the project and supervised the research. JJ, FC and DB developed the code; JJ performed the analysis. JJ, FC, and DB wrote the manuscript and approved the final version.

## Conflict of Interests

DB has a significant financial interest in GalaxyWorks, a company that may have a commercial interest in the results of this research and technology. This potential conflict of interest has been reviewed and is managed by the Cleveland Clinic.

JJ and FC have no conflicts to disclose.

## Funding

This work was supported by the Wellcome Trust [313498/Z/24/Z].

---

**Figure S1:**
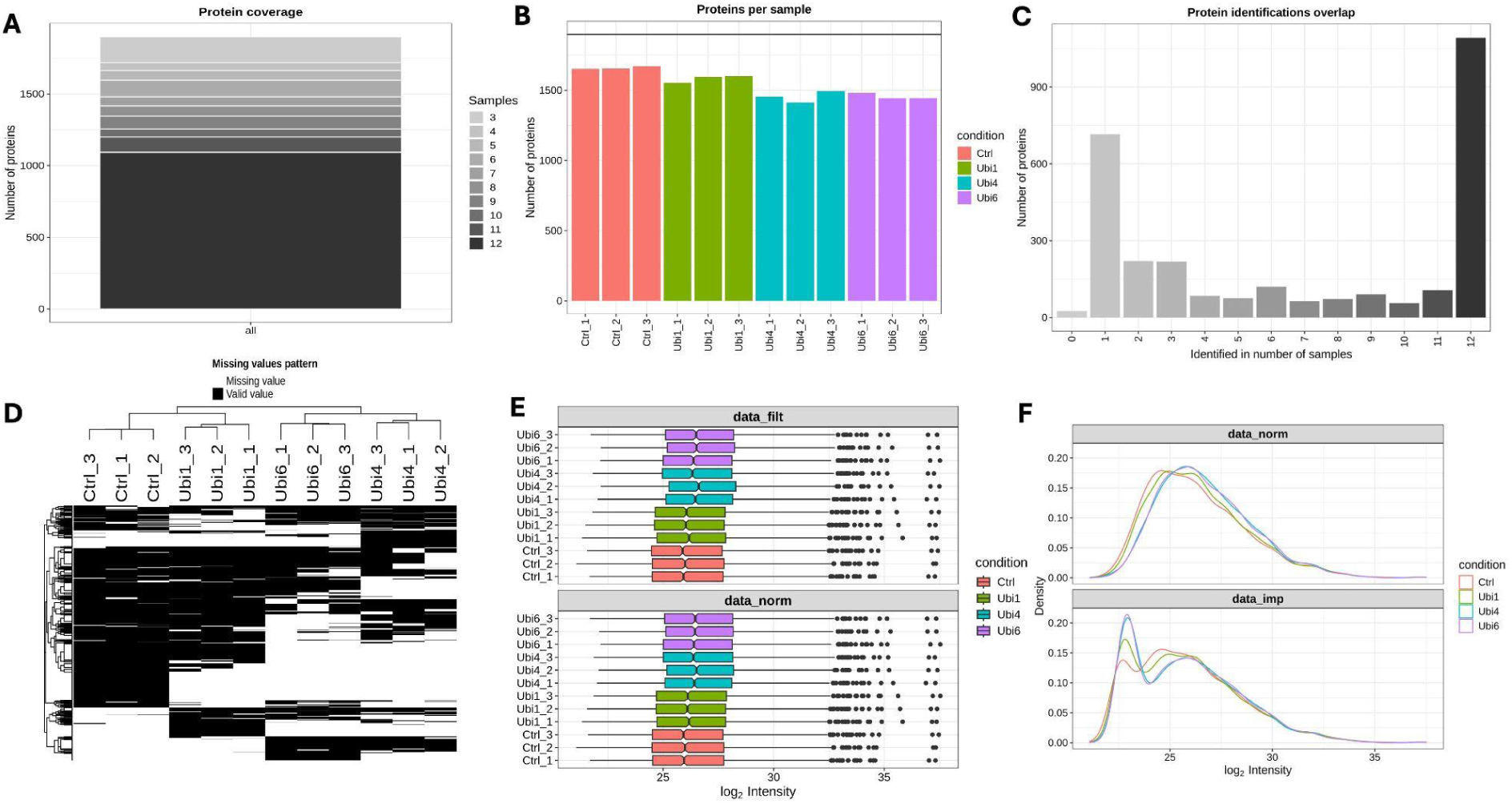
Figure demonstrates the automated pre-processing outputs generated by the ***DEP_pre_processing tool***, includes **(A)** *protein coverage*, **(B)** proteins per sample, **(C)** Protein identification overlap, **(D)** Missing value pattern heatmap, **(E)** Intensity distributions before and after normalization and **(F)** density plots of normalized and imputed data

**Figure S2:**
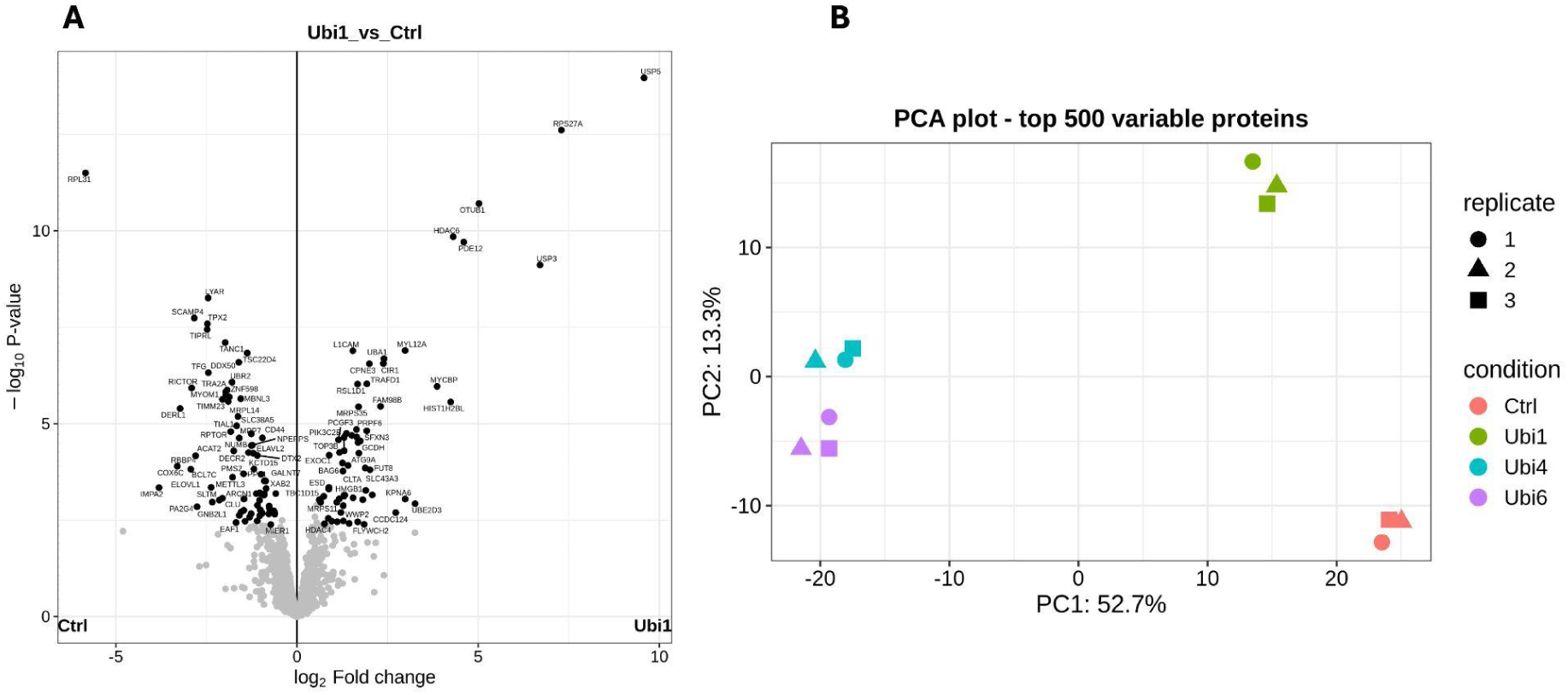
This figure highlights how the ***DEP_DE_analysis*** tool generates key downstream outputs, **(A)** differential expression and **(B)** PCA clustering, allowing users to interpret proteomic responses, verify sample grouping, and assess global expression changes across experimental conditions.

## Notes

https://github.com/BlankenbergLab/r2g2

https://github.com/jaidevjoshi83/galaxy_tool_wrappers

### Competing Interest Statement

Daniel Blankenberg has a significant financial interest in GalaxyWorks, a company that may have a commercial interest in the results of this research and technology. This potential conflict of interest has been reviewed and is managed by the Cleveland Clinic. Jayadev Joshi and Fabio Cumbo have no conflicts to disclose.

